# HIP-MS: An ultra-high-throughput, sensitive, and versatile affinity enrichment platform for static and dynamic interactome profiling

**DOI:** 10.64898/2026.06.03.729763

**Authors:** Louisa Grauvogel, Elisabeth Zollbrecht, Tim Heymann, Vincenth Brennsteiner, Christian Pensl, André C. Michaelis, Matthias Mann

**Author notes:** These authors contributed equally to this work (Louisa Grauvogel, Elisabeth Zollbrecht).

## Abstract

Although protein-protein interactions govern virtually all cellular processes, systematic interactome mapping by affinity enrichment mass spectrometry (AE-MS) is constrained by manual sample preparation and lengthy liquid chromatography (LC)-MS/MS acquisition. Here we present High-throughput Interactome Profiling by MS (HIP-MS), an automated, end-to-end pipeline that overcomes these limitations. It leverages the compact, high-affinity ALFA tag for on-plate nanobody capture in 384-well format, combined with on-plate tryptic digestion. It can process almost 10,000 samples per week from protein expression up to MS measurement and can be combined with ultra-fast gradient LC-MS acquisition of 500 samples per day. HIP-MS remains sensitive down to low-microgram lysate inputs, a 4,000-fold reduction compared to recent large-scale screens. Our pipeline recovers complexes from diverse cellular compartments and resolves endogenous membrane receptor signaling. HIP-MS establishes a scalable foundation for systematic interrogation of protein interactions across conditions, perturbations, and time, and for the generation of large-scale datasets for computational modeling.

## Introduction

Protein–protein interactions (PPIs) are fundamental to biology, with an estimated 80% of proteins functioning through interactions with other proteins^1^. Mass spectrometry (MS)-based proteomics has emerged as a central technology for systematically interrogating proteins and their interactions^2^. While complementary approaches such as yeast two-hybrid screening^3^ and proximity labeling^4^ have fundamentally contributed to PPI mapping, affinity purification mass spectrometry, or affinity enrichment MS (AE-MS) when performed quantitatively^5^, remains a gold standard for systematic interactome studies^5^. In AE-MS, protein complexes are captured under near-native conditions through an affinity tag fused to the bait and identified directly by MS, without the need for crosslinking, enzymatic labeling, or fluorescent reporters. AE-MS has historically had two main constraints: liquid chromatography (LC) throughput – i.e. samples Per Day (SPD) and the required input material per pulldown, which are both related to MS sensitivity and scan speed. Pioneering proteome-scale screens in yeast required liters of culture per bait, as limited enrichment efficiency and MS sensitivity drove milligram-scale input, while slow-scanning instruments and long nano-LC gradients restricted acquisition to a few samples per day^6,7^. Across subsequent applications spanning virus–host, human, and organelle-level interactomes, MS throughput increased more than 10-fold and input requirements likewise dropped by one to two orders of magnitude (Fig. 1a, b)^8–15^. However, even landmark efforts such as BioPlex still required years of instrument time at ∼20 SPD with milligram-level input^12,13,16^. A first comprehensive interactome study reported the entire yeast interactome in only ∼20 weeks by miniaturizing AE-MS to a 96-well plate format, measured at 60 interactomes per day^17^. That effort rested on two advances: a trapped ion mobility time-of-flight mass spectrometer (timsTOF) provided the sensitivity and scan speed for reduced input (350–500 µg per pulldown)^18^, while a chromatography system with disposable tips as trap columns and preformed gradients (Evosep One^19^) enabled 60 SPD via minimal overhead.

**Figure 1.**
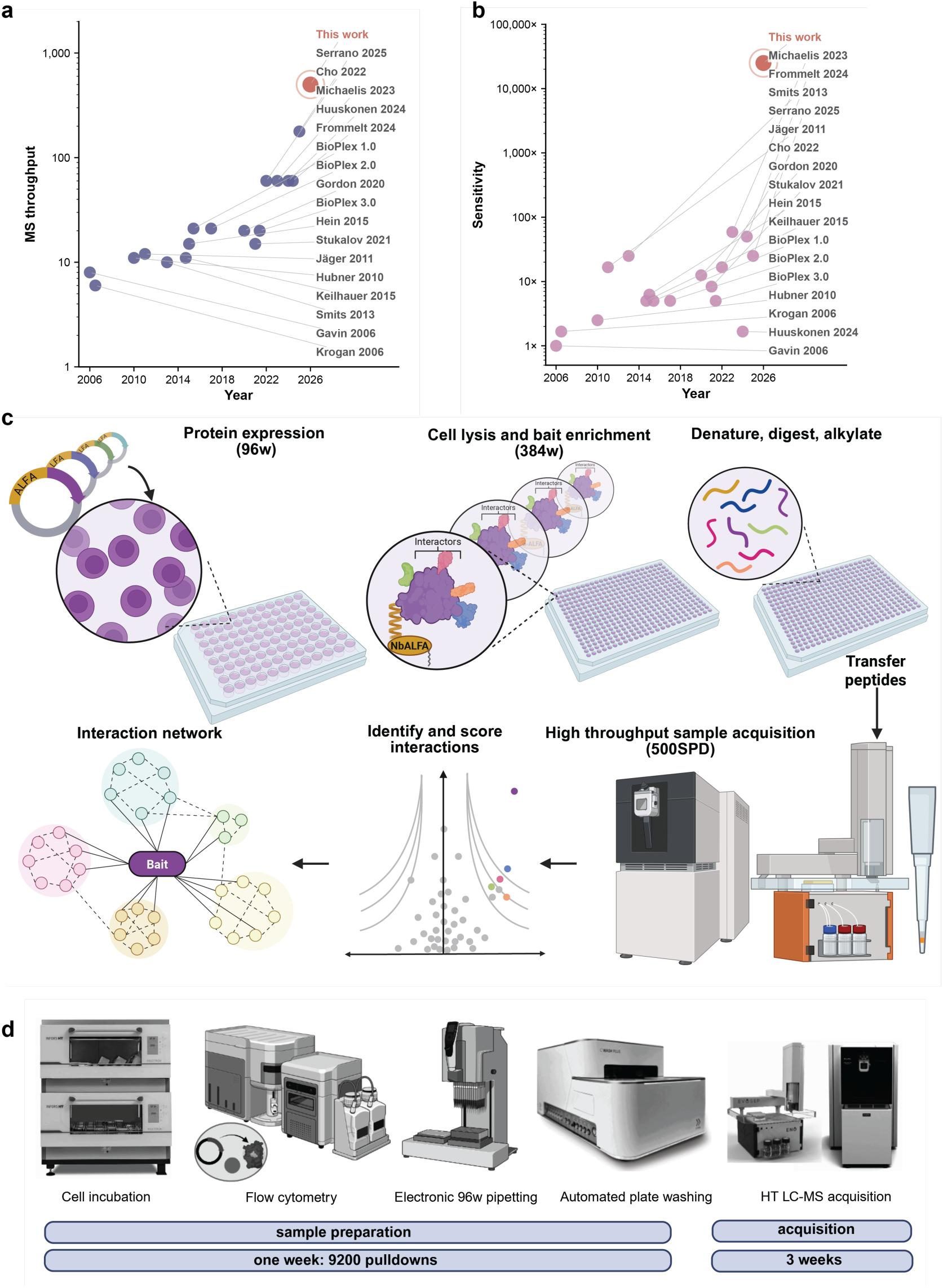
HIP-MS: an automated 384-well pipeline for high-throughput interactomics. (a) Mass-spectrometry acquisition throughput across published AE-MS / co-IP / TAP-MS interactome studies. y-axis: samples per day (SPD; calculated as 1,440 min / per-sample LC cycle time), log scale; x-axis: publication year. Each point is one study (purple); “This work” highlighted in coral at 500 samples per day (SPD). Studies plotted: Gavin et al.^6^, Krogan et al.^7^, Hubner et al.^25^, Jäger et al.^14^, Smits et al.^26^, Keilhauer et al.^5^, Hein et al.^10^, Huttlin et al. ^13^, Huttlin et al. ^16^, Gordon et al.^9^, Stukalov et al.^15^, Huttlin et al. ^12^, Cho et al.^8^, Michaelis et al.^17^, Huuskonen et al.^27^, Frommelt et al.^28^, Serrano et al.^21^. Throughput values were extracted from the methods section of each publication; where the LC cycle was not stated directly the SPD was estimated from the reported gradient length plus loading/ wash/ re-equilibration time. (b) Per-pulldown input requirement for the same set of studies, expressed as fold-reduction in starting protein mass relative to Gavin et al. 2006 (arbitrarily set to 1×; log scale). Input mass refers to the total protein loaded onto bait-bound resin in a single pulldown; for studies that fractionated samples post-pulldown the value reflects pre-fractionation input (c) Cartoon of the HIP-MS workflow. ALFA-tagged ORFs are transiently transfected into cell lines in 96-well format, natively lysed and transferred to 384-well plates pre-coated with the NbALFA nanobody, incubated for 3h (or overnight) and digested directly on the plate. The resulting peptides are analyzed on an Orbitrap Astral Zoom coupled to an Evosep Eno liquid chromatography system and measured at 500 SPD. Schematic created in BioRender. (d) Hardware and timeline of the HIP-MS pipeline: cell incubation (CO2 incubator), flow-cytometry expression QC, electronic 96-well pipetting, automated 384-well plate washer, and the Orbitrap Astral Zoom with Evosep Eno.

The recent introduction of a hybrid Orbitrap–time-of-flight instrument (Orbitrap Astral) allowed even faster and deeper proteome coverage and has already been employed for very fast interaction measurements^20–22^. In parallel, the Evosep Eno further improved chromatographic peak shape in gradients of ∼2.9 min per sample, for 500 SPD throughput^23^.

While instrumentation advances have steadily eased the analytical bottleneck, the upstream sample preparation side of AE-MS has not kept pace. The 96-well workflow described above still requires about half a milligram of lysate per pulldown and is largely manual^17^. Most other larger-scale studies operate even in the milligram range, requiring multiple 10-cm dishes per bait, manual bead handling, and stable cell lines generated by lentiviral transduction^12^. This combination of experimental steps takes months per construct and is difficult to scale beyond a few hundred baits. Transient transfection would be attractive because it can be performed in 12–48 h from a 96-well plasmid plate, scaling to hundreds of baits in parallel and can be ported across cell lines without re-engineering. However, the per-well yield falls well below the milligram-scale input convention AE-MS requires. Together, these constraints become especially limiting for weakly expressed proteins, membrane receptors, and time-resolved experiments, where input material is intrinsically scarce or where the number of conditions multiplies the per-experiment cost. Addressing this gap requires a workflow that simultaneously reduces per-bait input, automates sample handling, and enables high-throughput MS acquisition.

Here we present High-throughput Interactome Profiling (HIP-MS), an automated, 384-well AE-MS pipeline that reaches 500 SPD acquisition ∼10,000 pulldowns per week of upstream sample preparation. By combining miniaturized input, automated on-plate capture, and transient expression in a single workflow, HIP-MS enables interactome experiments that have previously been out of reach, including endogenous membrane receptor signaling captured with temporal resolution and systematic perturbation screens across many baits in parallel.

## Results

### HIP-MS: an automated 384-well pipeline for high-throughput interactomics

To transform interactomics into a systems-level approach, in which thousands of baits can be screened across conditions, perturbations, and cell lines, we developed HIP-MS, an automated, 384-well, on-plate AE-MS pipeline capable of processing ∼9,200 pulldowns per week **(Fig. 1c, d).** The platform brings together five components, each addressing a distinct challenge of AE-MS workflows: the affinity tag, the bait expression strategy, the cell culture and lysis format, the enrichment and washing chemistry, and the LC-MS acquisition. HIP-MS integrates these parts into a single end-to-end workflow operating at high throughput.

#### Affinity tag

Choosing the tag system for AE-MS involves three competing requirements: binding tight enough to survive washing steps, being small enough to avoid perturbing the bait’s function and localization and contributing minimal background from the capture reagent itself. We use the ALFA-tag system as it satisfies all three requirements^24^. The ALFA-tag itself is only 15 amino acids long (1.7 kDa) and by design folds into a defined alpha-helical structure which minimizes the risk of steric interference or functional perturbation of the bait, a recurring concern with larger tags such as GFP or tandem FLAG-HA used in previous proteome-scale screens^10,12^ **(Supplementary Fig. 1a)**. Its cognate nanobody (NbALFA) binds with picomolar affinity, considerably stronger than common tag systems **(Supplementary Fig. 1b).** As we digest directly on the plate without an elution step **(Fig. 1c)**, this high affinity is a pure advantage, as it avoids the binding affinity trade-off of conventional, elution-based AE-MS workflows. NbALFA itself is likewise compact (∼13 kDa) and yields only 10 MS-visible tryptic peptides, limiting the contribution of the capture reagent itself to the MS background– circumventing a major contamination problem observed with the use of antibodies based approaches.

#### Bait expression

Generating a stable cell line for each bait by lentiviral transduction or genome editing takes months and is one of the rate-limiting steps in conventional large-scale interactome screens. HIP-MS instead uses transient transfection, allowing hundreds of constructs to be expressed in parallel within 12–24 h. As constructs only need to be delivered to cells rather than integrated into the genome, the same workflow ports directly across cell lines without re-engineering. Cell viability and transfection efficiency can be monitored throughout by flow cytometry (suspension cells) or live-cell imaging (adherent cells) **(Fig. 1d).**

#### Cell culture and native lysis

Capturing endogenous interactions requires preserving transient and weak associations that are easily disrupted by denaturing buffers or elevated temperatures. We therefore carry out cell culture, transfection, and native lysis in 96-well format using portable 96-channel pipetting workstations that can operate at both room temperature and 4 °C inside a cold room **(Fig. 1d).** We optimized the lysis chemistry (an IGEPAL-based buffer supplemented with benzonase) for complex preservation; benzonase clears nucleic acids to reduce viscosity for automated handling in 96-well format (see Methods). We then transfer lysates to 384-well plates for affinity enrichment, quadrupling the number of samples per plate **(Fig. 1c).**

#### On-plate enrichment and washing

In our experience, manual bead handling is the largest source of variability and labor in conventional AE-MS, and the associated sample losses scale poorly with input amount. HIP-MS instead applies native lysates to 384-well plates coated with streptavidin-immobilized NbALFA, capturing bait–prey complexes directly on the plate surface. Compared with bead-based capture, the plate format presents far less surface area for non-specific binding and allows deterministic wash cycles with complete buffer exchange. Automated plate washing removes non-specific background and detergent, and captured complexes are then digested and alkylated directly on plate without a separate cleanup step **(Fig. 1c, d)**. Because every well receives an identical, programmable number of wash cycles, the background proteome remains consistent across samples within a batch. This is essential for statistical comparisons across pulldowns and can be tuned to the experiment, with more stringent washing favoring stable complexes and less stringent washing preserving transient interactions. The on-plate workflow eliminates bead handling entirely and avoids the sample losses associated with multiple transfer steps, bringing input requirements down to the low microgram range, a ∼4,000-fold reduction relative to recent large-scale interaction studies^12^ **(Fig. 1b**, **Fig. 4a).**

#### LC-MS acquisition

The throughput of AE-MS has historically been limited by the speed of MS acquisition and the total time needed for each LC run. We address this by coupling an Orbitrap Astral Zoom mass spectrometer to an Evosep Eno LC system, acquiring pulldowns at 500 SPD. We found that the Astral Zoom provides the sensitivity and scan speed required to acquire interaction samples at short gradients, while the Evosep Eno contributed sharp peaks and minimized cycle overhead, together making 500 SPD robustly accessible for AE-MS. This represents a ∼25-fold increase over proteome-scale human interactome studies performed at conventional nano-flow LC^12^, a ∼3-fold increase over the fastest current AE-MS methods^21^, and a >60-fold increase relative to the first proteome-scale AE-MS screen^6^ **(Fig. 1a).**

### Shorter gradients preserve the core interactome

To benchmark our pipeline, we selected 25 representative bait proteins spanning multiple subcellular compartments: soluble cytosolic and nuclear baits, plus six membrane-associated baits (EMC2, EMC3, COX5B, MICU1, OSTC, BECN1) that are typically more challenging to capture **(Fig. 2a).** We designed each bait as a C-terminal ALFA-tagged plasmid construct and transiently expressed the tagged proteins in HEK cells.

**Figure 2.**
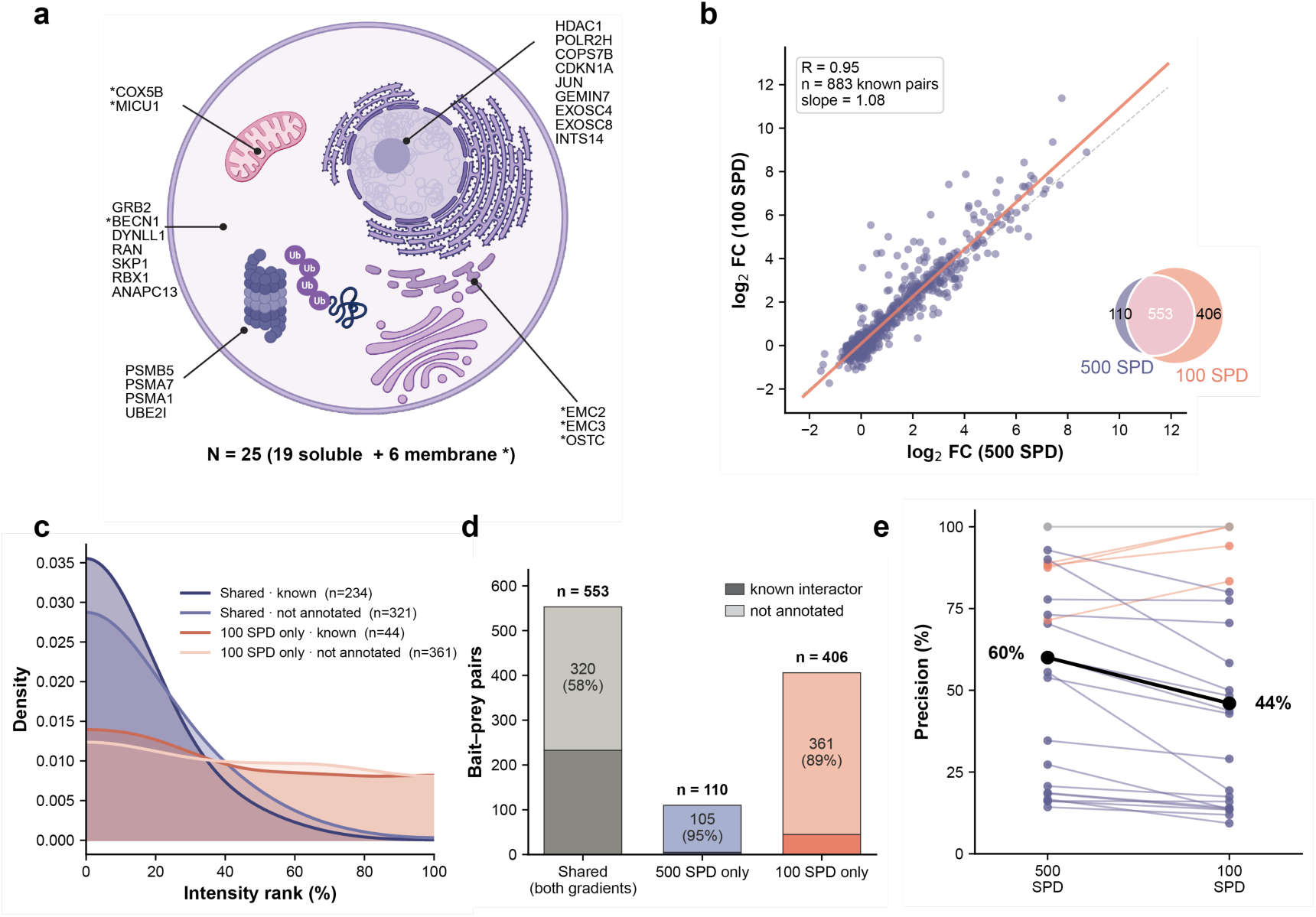
Shorter gradients preserve the core interactome. (a) Cell-compartment schematic of the 25 curated baits with their subcellular locations (UniProt-derived membrane annotation; soluble =unmarked, membrane-encoded marked with an asterisk). Cartoon created with BioRender. (b) Log₂ fold-change agreement across gradients for known bait–prey pairs. Each dot is one bait (STRING / Complex-Portal-annotated interactor pair). Inset: overlap of significantly enriched bait–prey pairs called at each gradient (Euler/Venn). (c) Kernel-density estimate of per-bait intensity-rank percentile (0 = highest abundance, 100 = lowest) for enriched bait–prey pairs, split by gradient agreement × STRING/Complex-Portal annotation. (d) Composition of enriched bait–prey pairs by gradient agreement (shared / 500-SPD only / 100-SPD only); each bar split into known interactors (dark) vs not annotated (light), with the not-annotated count and percentage labelled and total n shown above each bar. (e) Per-bait precision at each gradient. Precision = the fraction of a bait’s significantly enriched proteins (FDR < 0.05, log₂ FC > 1.5) that are STRING / Complex-Portal-annotated interactors of that bait. Bold black line represents the medians (n = 24 baits, COX5B excluded).

Before characterizing the recovered interactomes in depth, we asked how far MS acquisition could be accelerated without losing biological signal, since acquisition speed sets the ceiling on screening throughput. We therefore analyzed the same 25 baits at 500 and 100 SPD on the Orbitrap Astral Zoom (2.3 min vs 13 min gradient time, respectively). Fold changes of known interactors were highly concordant between gradients (Pearson R = 0.95, n = 883 known bait–prey pairs; **Fig. 2b**), far exceeding the cross-gradient correlation across all detected pairs (R = 0.70, n = 154,475; **Supplementary Fig. 2a,b**). This indicates that enrichment strength is maintained at five-fold higher MS throughput and that the high reproducibility reflects genuine biological interactions rather than a technical baseline alone.

As expected, the 100 SPD method identified more proteins per pulldown overall (median ∼3,700 vs ∼7,000; 1.9-fold deeper) **(Supplementary Fig. 4a)**. However, the two gradients agreed on the large majority of differentially enriched bait–prey calls (553 shared), with 110 unique to 500 SPD and 406 unique to 100 SPD **(Fig. 2b, inset).** To investigate whether the additional depth gained with the longer gradient increases biological signal for interactors, we asked where along the intensity distribution the additional differentially enriched proteins arise. We ranked significantly upregulated proteins by their per-bait intensity percentile and classified them as shared with the 500 SPD dataset or unique to 100 SPD, further split by whether they corresponded to a known interactor (STRING combined score > 700 or Complex Portal) **(Fig. 2c).** Shared hits concentrated at the top of the intensity distribution and were dominated by known interactors. Differences emerged primarily at the lower-intensity end of the distribution: hits unique to 100 SPD extended much further into the low-abundance range, where non-annotated proteins outnumbered known interactors roughly 8:1 (361 vs 45 pairs) **(Fig. 2c).** These additional identifications likely included some substoichiometric or transient interactions, but as shown below, they were dominated by un-annotated proteins. To quantify the biological yield of each gradient, we partitioned all DE bait–prey pairs across the 25 curated baits into three categories: shared (DE in both gradients), 500 SPD only, and 100 SPD only, and asked what fraction of each corresponded to a known interactor. Shared pairs were 3.8-fold enriched for known interactors (42.1% of 553 pairs) compared with pairs unique to 100 SPD (11.1% of 406), and 9.3-fold enriched compared to pairs unique to 500 SPD (4.5% of 110) **(Fig. 2d).** Consistent with this, the per-bait precision, defined as the fraction of significantly enriched proteins corresponding to an annotated interactor (STRING combined score > 700 or Complex Portal, FDR < 0.05, log₂ FC > 1.5) was higher at 500 SPD than at 100 SPD (median 60% vs 44%; higher at 500 SPD in 18 of 24 baits enriched in both gradients; **Fig. 2e**). Thus, although the longer gradient extended detection into lower-abundance proteins and recovered some additional interactors, its extra identifications were dominated by un-annotated, low-intensity pairs suggesting that the added depth comes at the cost of precision.

Together, these findings show that 500 SPD preserves the high-confidence interactome at five-fold higher MS throughput while reducing unspecific background.

### Recovery of protein complexes across cellular compartments

Having established that 500 SPD preserved the high-confidence interactome, we characterized the recovered complexes at this throughput for our 25-bait panel. In each pulldown, we robustly enriched the bait together with its known interaction partners (STRING and Complex Portal), ranking among the most strongly enriched proteins. The representative baits spanned the major subcellular compartments: PSMB5 (20S proteasome) in the cytosol, POLR2H (RNA polymerase II) in the nucleus, EXOSC4 (RNA exosome), EMC2 (ER-membrane EMC complex), COX5B (cytochrome c oxidase) in the mitochondria, and RBX1 (cullin-RING ubiquitin ligase) **(Fig. 3a).** In each pulldown, the bait and its established interactors ranked among the most strongly enriched proteins, showing consistent recovery across diverse subcellular environments. For the membrane-resident COX5B pulldown, four subunits of the cytochrome c oxidase complex (COX4I1, COX5A, COX6B1, MT-CO2) co-enriched with the bait at strong significance (FDR < 10^-4^) despite modest fold-changes (log_2_ FC = 1.1-1.2), demonstrating that our pipeline captures tightly bound membrane-embedded interactors **(Fig. 3a)**.

**Figure 3.**
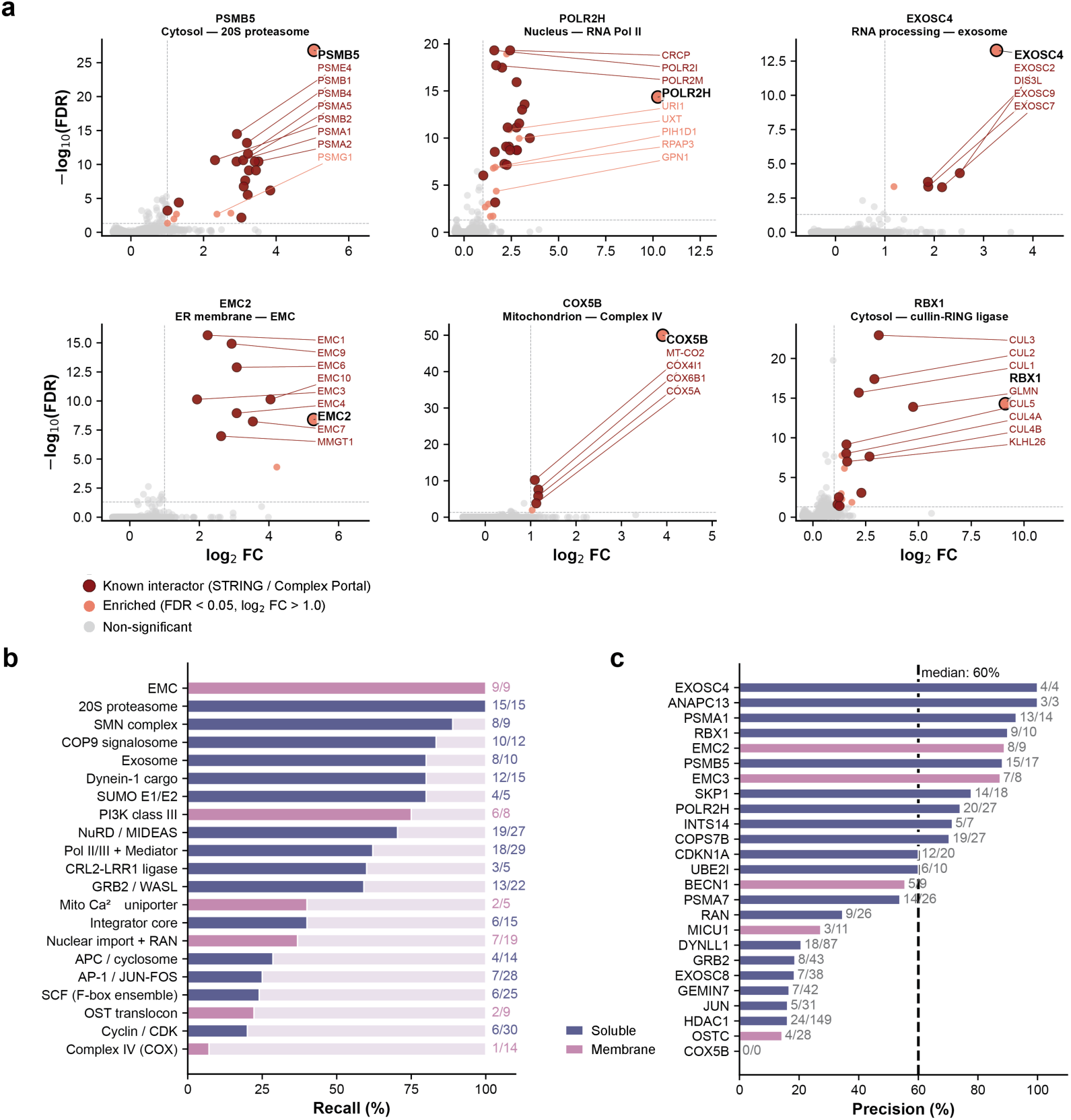
Recovery of protein complexes across cellular compartments. (a) Volcano plots for six representative baits spanning subcellular compartments: PSMB5 (cytosol, 20S proteasome), POLR2H (nucleus, RNA PolII), EXOSC4 (RNA processing, exosome), EMC2 (ER membrane, EMC), COX5B (mitochondrion, Complex IV), and RBX1 (cytosol, cullin-RING ubiquitin ligase). Each point is one protein group. FDR < 0.05, log₂ FC > 1.0. (b) Recall of protein-complex subunits across 21 Complex Portal complexes. For each curated bait, the highest-recovered CP complex of size 5–30 was selected automatically (with a manual size cap to exclude CP “umbrella” entries that lump many sub-complexes together); when multiple baits land on the same CP complex (e.g., PSMB5, PSMA7, PSMA1, 20S proteasome; EMC2, EMC3, EMC), they collapse to one row. X-axis = % of CP subunits recovered at FDR < 0.05, log₂ FC > 1.5. (c) Per-bait precision: fraction of significantly enriched proteins (FDR < 0.05, log₂ FC > 1.5) annotated as known STRING / Complex Portal interactors.

To assess complex recovery (recall), we mapped each curated bait to its best-matching Complex Portal entry, yielding 21 unique complexes, covering all 25 baits **(Fig. 3b).** Across these complexes we recovered 166 of 325 annotated subunits (51%). This is a deliberately conservative lower bound: it counts every annotated subunit of every complex, including large assemblies probed through only a single, peripheral entry-point bait. Because HIP-MS routinely screens many baits in parallel, most complexes will in practice be probed from multiple subunits, causing recovery to rise sharply. The most extreme example is Complex IV, where only 1 of its 14 subunits passed the stringent fold-change threshold from the single COX5B pulldown, even though additional cytochrome c oxidase subunits were detected at lower fold-change **(Fig. 3a)**. Complexes captured through their core subunits were almost recovered completely: the EMC (9/9; EMC2 + EMC3) and the 20S proteasome (15/15; PSMB5 + PSMA7 + PSMA1) reached 100%, with ≥80% recovery for the SMN complex (8/9), COP9 signalosome (10/12), cytoplasmic exosome (8/10), dynein-1 cargo (12/15), and SUMO E1/E2 conjugation (4/5). The high per-complex recovery achieved from individual bait pulldowns therefore indicates that the 51% aggregate substantially underestimates the assay’s sensitivity.

To quantify precision, we calculated the fraction of significantly enriched proteins annotated as known interactors in STRING (combined score > 700) or Complex Portal **(Fig. 3c).** The median across the 25 baits was 60%, with ANAPC13 (3/3) and EXOSC4 (4/4) reaching 100%, and PSMA1 (93%), RBX1 (90 %), EMC2 (89%), and PSMB5 (88%) above 85%. Baits with the highest precision tended to recover fewer co-enriched proteins overall, whereas baits with broader interaction landscapes (HDAC1, DYNLL1) pulled down a larger fraction of unannotated proteins, consistent with intrinsic interaction promiscuity rather than data quality driving this trend.

These precision values represent conservative lower bounds because of incomplete database coverage. For example, proteins with well-established roles at the bait’s complex were strongly enriched yet not annotated as interactors in either database, e.g. the 20S-proteasome assembly chaperone PSMG1 in the PSMB5 pulldown (log₂ FC = 2.4, FDR = 2×10⁻³), and several subunits of the R2TP/PAQosome co-chaperone that mediates RNA polymerase II assembly in the POLR2H pulldown (URI1, log₂ FC= 2.4, FDR = 1×10⁻¹¹; UXT, log₂ FC = 2.9; RPAP3; PIH1D1; GPN1) **(Fig. 3a)**. HIP-MS thus recovers complexes with high precision across every major subcellular compartment, from soluble cytosolic and nuclear assemblies to membrane-embedded mitochondrial and ER complexes.

### Sensitivity and versatility across cell lines

Many biologically interesting systems are intrinsically sample-limited: rare cell populations, hard-to-transfect cells, and endogenously low-abundance proteins all yield far less material than conventional AE-MS requires. Thus, scaling down input is a prerequisite for extending interactome mapping beyond high-yielding cell lines. We titrated the input for EMC2-ALFA pulldowns across more than three orders of magnitude, from 100 µg down to 39 ng of HEK293T cell lysate **(Fig. 4a)**. While proteome depth scaled with input, dropping from a median of 4,411 to 99 quantified proteins per sample across this range, complex enrichment was retained at lower inputs **(Supplementary Fig. 4d)**. At 50–100 µg of lysate, eight EMC subunits were significantly enriched, recovering essentially the entire complex. At only 1.25 µg, we still found four EMC subunits enriched: EMC2, EMC3, EMC4, and EMC8. The bait itself was still robustly captured at just 312 ng of lysate, suggesting that we can specifically enrich targets from sub-microgram input material, more than three orders of magnitude below the milligram-scale material used in proteome-scale AE-MS screens.

To investigate how our workflow translates across cell lines, we applied it to adherent HEK293T cells in 96-well format, corresponding to only ∼10 µg of input lysate per pulldown. Despite the much lower background proteome of the adherent cells (median 920 vs 3,756 protein identifications; **Supplementary Fig. 4a**), per-bait precision and recall were comparable between the two cell lines (median precision 44% in 293T vs 60% in 293F at 500 SPD; median recall 50% vs 60%), indicating that the lower proteome depth in adherent cells did not substantially compromise biological signal **(Fig. 4b)**. Fold changes of known interactors at 500 SPD were well correlated between suspension and adhesion cells (Pearson R = 0.73 across 630 bait-interactor pairs spanning 25 curated baits; **Fig. 4c**), with a regression slope of 0.79 indicating a mild compression of the dynamic range in 293T. Note that this correlation was significantly stronger than the baseline correlation across all detected bait–prey pairs (R = 0.34, n = 82,650 pairs), confirming that the cross-cell-line reproducibility is driven by genuine biological interactions and not by shared technical baseline **(Supplementary Fig. 4b,c)**.

At the level of protein complexes, completeness was largely preserved across the two cell lines **(Fig. 4d).** Core complexes such as the 20S proteasome (15 of 15 in 293F, 14 of 15 in 293T), the EMC (9 of 9 in 293F, 8 of 9 in 293T), and the COP9 signalosome (10 of 12 vs 7 of 12) reached near-complete recovery. Notably, several membrane complexes including Complex IV (cytochrome c oxidase, 6 of 14 subunits in 293T vs 1 of 14 in 293F) and the OST translocon (5 of 9 vs 2 of 9) recovered better in 293T.

**Figure 4.**
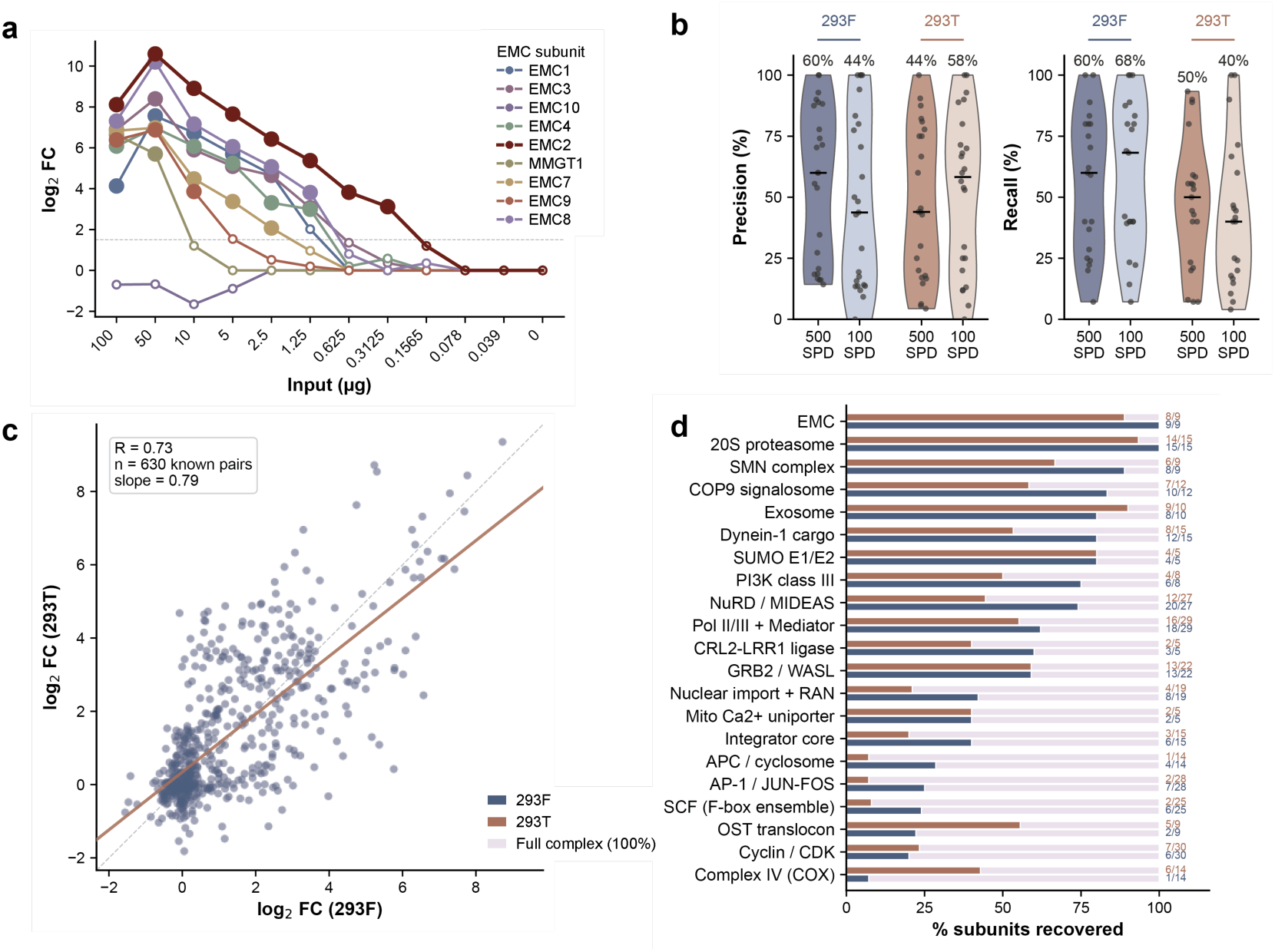
Sensitivity and versatility across cell lines. (a) EMC2 input titration in HEK293T cells (500 SPD). Enrichment (log_2_ fold-change vs GFP) of EMC subunits across an input series from 0 to 100 µg of lysate. Each line is one EMC subunit. Hollow circles denote detected but non-significant proteins (FDR < 0.05, log_2_ FC > 1.5). (b) Per-bait precision and per-complex recall across both cell lines and gradients (293F and 293T at 500 and 100 SPD). Precision is computed per bait as the fraction of significantly enriched proteins (FDR < 0.05, log_2_ FC > 1.5) that are annotated interactors of that bait in STRING or Complex Portal, with the bait itself excluded from both numerator and denominator. Recall: for each of 21 Complex Portal complexes, the union of subunits recovered across the baits targeting that complex divided by the complex size. (c) Cross-cell-line reproducibility of enrichment. Log_2_ fold-changes of annotated interactors (STRING or Complex Portal) measured in 293F (x) versus 293T (y) at 500 SPD (d) Complex completeness in 293F versus 293T at 500 SPD. For each of the 21 Complex Portal complexes, bars give the percentage of subunits recovered (enriched in at least one targeting bait); paired bars compare 293F (upper) and 293T (lower).

Thus our workflow generalizes to a second, adherent cell line and to inputs up to ∼4,000-fold below the conventional scale, making interactome mapping feasible for diverse cell types and sample-limited systems.

### Endogenous TNFR1 signaling captured with temporal resolution

To demonstrate the biological utility of our pipeline beyond steady-state interactomes, we applied it to two dynamic systems with well-characterized interaction landscapes: TNF receptor signaling and rapamycin-induced autophagy.

Membrane receptors are a particularly challenging class of baits for AE-MS. They typically have low copy numbers, partition into detergent-sensitive membrane environments, and engage cytoplasmic adaptors transiently and substoichiometrically^29^. As a result, prior characterizations of the TNFR1 signaling complex have generally relied on either receptor overexpression or large-scale input from large-format adherent cultures^30^. To test whether our miniaturized platform can capture an unmodified, endogenously expressed membrane receptor and its intracellular signaling complex in a single 96-well experiment, we used an ALFA-tagged, isoleucine zipper-stabilized TNF cytokine (ALFA-IZ-TNFα) as an extracellular bait and confirmed activation of the canonical TNF signaling cascade by phosphoproteomics **(Fig. 5a, Supplementary Fig. 5b and 5c).** Importantly, TNFR1 itself was not overexpressed. Instead, we captured it directly from the endogenous receptor pool of A549 cells via the extracellular ligand.

**Figure 5.**
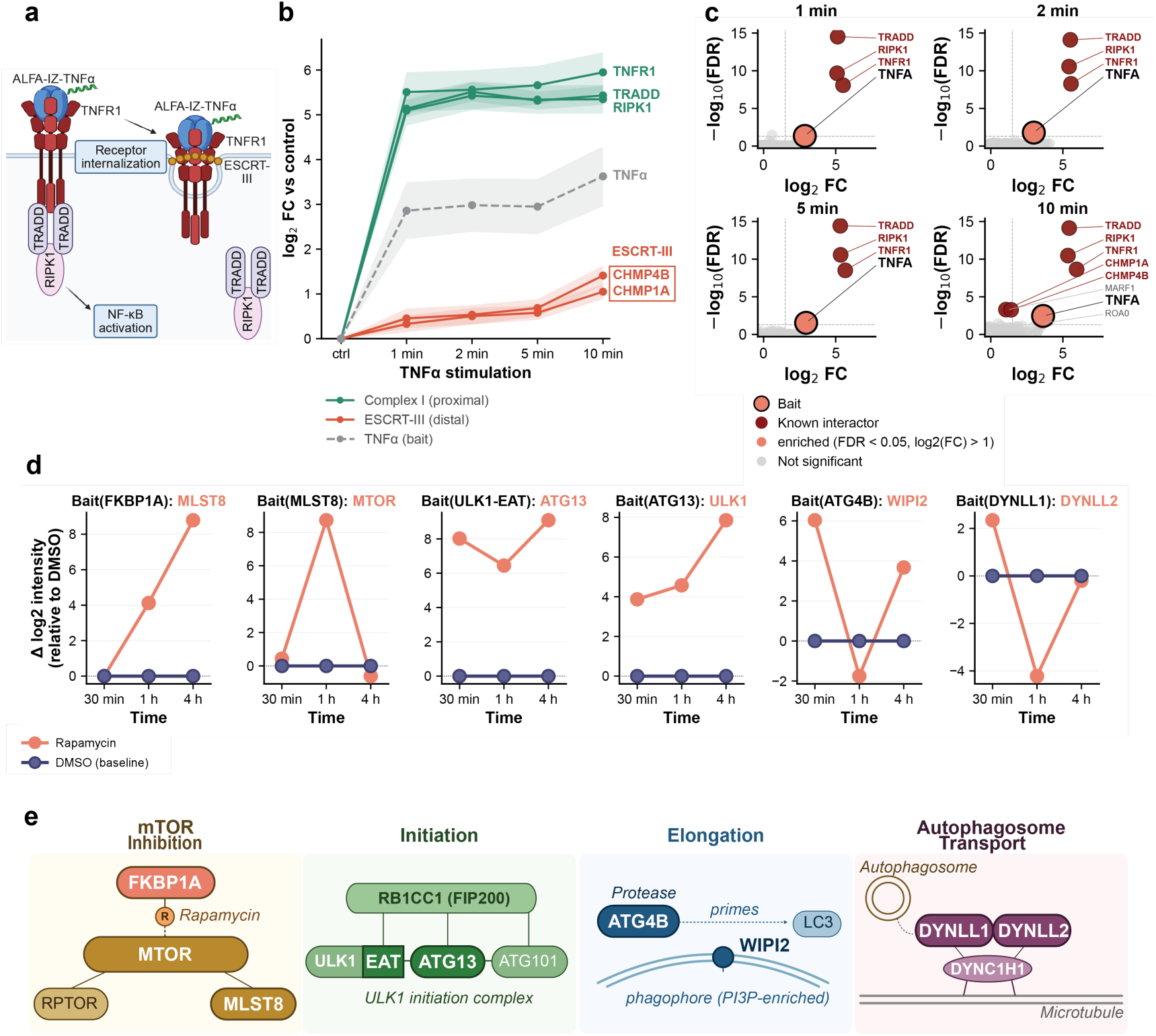
Time-resolved interactome mapping of TNF signaling and rapamycin-induced autophagy. (a) Schematic of TNFα–TNFR1 signaling. We use an ALFA-tagged isoleucine-zipper TNFα (ALFA-IZ-TNFα), enabling capture of endogenous TNFR1 and downstream effectors. (b) Mean log₂ fold-change versus unstimulated control for Complex I components (TNFR1, TRADD, RIPK1; green), ESCRT-III subunits (CHMP1A, CHMP4B; red), and bait (TNFα; gray dashed) in ALFA-IZ-TNFα pulldowns at 1, 2, 5, and 10 min of TNFα stimulation. Shaded bands: ±1 SEM (n = 12 per timepoint). A549 cells, 100 SPD. (c) Volcano plots for the same pulldowns (four stimulation timepoints) versus unstimulated control. Dark red: known interactors (STRING / Complex Portal); black outline: bait; dashed lines: |log₂FC| > 1, FDR < 0.05. (d) Median Δ log₂ intensity (Rapamycin-treated minus vehicle) for six baits and three treatment timepoints (30 min, 1 h, 4 h). In each subpanel the DMSO trace is pinned at zero by definition. (e) Schematics of the four autophagy stages probed in d. *mTOR inhibition*: FKBP1A–rapamycin docking onto MTOR within mTORC1, alongside MLST8 and RPTOR. *Initiation*: the RB1CC1 (FIP200)-scaffolded ULK1 initiation complex with the ULK1 EAT subdomain bound to ATG13 and ATG101. *Elongation*: ATG4B priming LC3, WIPI2 recruited to the PI3P-enriched phagophore membrane. *Autophagosome transport*: DYNLL1/2 with DYNC1H1 on microtubules.

Pulldowns at four stimulation timepoints (1, 2, 5, and 10 min; n = 12 per condition) revealed two temporally separated states of the TNFR1 signaling cascade **(Fig. 5b, c).** The canonical plasma-membrane Complex I — TNFR1 together with the cytoplasmic adaptors TRADD and RIPK1 — was significantly co-enriched within 1 min and remained stably bound across the time-course (log_2_ FC ≈ 5-6, FDR < 10−8 at every timepoint; Fig. 5c). By 10 min we additionally detected the ESCRT-III components CHMP1A and CHMP4B (log_2_ FC ≈ 1.0-1.4), consistent with receptor internalization and endosomal sorting following ligand engagement. The two modules were kinetically separable: Complex I plateaued within 1 min, whereas ESCRT-III became prominent from ∼5 min onwards **(Fig. 5b)**. Thus, HIP-MS co-captured the membrane-embedded receptor and its cytoplasmic signaling adaptors from the same lysate, recovering both the receptor-proximal and the internalization-associated arms of the complex. Other canonical Complex I members including TRAF2, cIAP1/2, and the LUBAC subunits were not significantly enriched across the time-course, likely reflecting their substoichiometric and transient association with the receptor.

To assess whether the miniaturized 96-well capture loses signal relative to a conventional workflow, we performed a side-by-side bead-based pulldown using ALFA Selector ST magnetic agarose beads from 10-cm dishes at 15 min of TNFα stimulation **(Supplementary Fig. 5d).** The bead format did not yield additional canonical TNFR1 interactors: TRAF2, cIAP1/2, and LUBAC subunits remained below significance, and TRADD, RIPK1, CHMP1A and CHMP4B were detected but did not reach the formal threshold at this single late timepoint. Importantly, the bead pulldown returned a much larger set of significantly enriched proteins than the plate format, the majority of which lack annotated TNF-signaling relevance and likely correspond to non-specific background associated with the larger surface area and input scale of the bead workflow. The miniaturized 96-well format therefore recovered the receptor-proximal interactome as completely as the conventional bead-based workflow while strongly reducing non-specific background.

### Stage-resolved mapping of rapamycin-induced autophagy

We next took advantage of the throughput and sensitivity of our platform to sample distinct stages of the same pathway in parallel. We expressed 23 ALFA-tagged constructs (12 autophagy genes, most in both N- and C-terminal tag orientations) spanning sequential stages of autophagy — mTOR inhibition, initiation, nucleation, elongation, autophagosome transport, and closure — in HEK293T cells and pulled them down at three rapamycin timepoints (30 min, 1 h, 4 h) alongside matched DMSO controls. Engagement of the mTOR/autophagy network at the pathway level was confirmed by parallel phosphoproteomics on the same cell system **(Supplementary Fig. 5e, f).**

FKBP1A, MLST8, ULK1-EAT, ATG13, ATG4B, and DYNLL1 captured canonical pathway partners with stage-consistent temporal behavior, resolving four sequential autophagy stages (mTOR inhibition, ULK1-complex initiation, elongation, and autophagosome transport) **(Fig. 5d, e).** Their drug-induced trajectories, expressed as the Δ log₂ intensity relative to matched DMSO controls, are summarized in **Fig. 5d**, with the corresponding enrichment significance in **Supplementary Fig. 5g**. The remaining bait constructs reproducibly expressed and immunoprecipitated their own bait protein but did not reach robust partner enrichment at 500 SPD, consistent with autophagy interactions being predominantly transient and substoichiometric, making this panel a stress test at the sensitivity edge of the current HIP-MS workflow.

FKBP1A captured MLST8 in a strictly rapamycin-dependent and time-progressive manner, with Δ log₂ intensity rising from ∼ 0 at 30 min to ∼ 9 at 4 h (significant from 1 h onward; combined p < 0.05 and |Δlog₂FC| > 1, **Supplementary Fig. 5g**). MTOR itself was below the protein-group quantification threshold in FKBP1A pulldowns, so MLST8 served as the proxy mTORC1 readout. Reciprocally, MLST8 recovered MTOR robustly in both rapamycin and DMSO conditions at every timepoint, identifying the mTORC1 core as a constitutive assembly that remains intact under rapamycin’s allosteric inhibition. Unlike the rapamycin-induced interactions, this recovery showed no time-progressive, rapamycin-dependent gain; the residual differences were small and non-monotonic (**Supplementary Fig. 5g**), with the 1 h excursion attributable to reduced MTOR detection in that plate rather than a change in complex occupancy.

ULK1-EAT captured ATG13 specifically under rapamycin at all three timepoints, consistent with the canonical model in which mTORC1 inhibition leads to ATG13 dephosphorylation and tighter binding to the ULK1 EAT subdomain. Critically, we independently observed the reciprocal interaction in the ATG13-ALFA pulldown, where ULK1 was likewise stably recovered under rapamycin treatment, providing orthogonal validation that the rapamycin-induced stabilization of the ULK1 initiation complex is captured from both sides of the interaction. ATG4B captured WIPI2 with positive Δ log₂ intensity at 30 min and 4 h, consistent with elongation-stage engagement of the ATG4B-LC3 conjugation machinery at the phagophore membrane. The transient drop at 1 h likely reflects imputation noise at that plate rather than a biological reversal, and the interaction was non-reciprocal in our dataset (no ATG4B recovery in WIPI2-ALFA pulldowns) and so should be regarded as a candidate finding pending orthogonal validation. Finally, DYNLL1 recovered its paralog DYNLL2 robustly in both rapamycin and DMSO conditions, consistent with the basal dynein light-chain heterodimer; as for MLST8–MTOR, this constitutive interaction showed no time-progressive rapamycin-dependent change.

Together, these results demonstrate that the platform resolves ligand-induced membrane receptor complex assembly and small-molecule-driven intracellular pathway remodeling within a single 96-well capture format, with temporal resolution sufficient to separate sequential complex states.

## Discussion

Experimental interaction mapping has remained fundamentally low-throughput relative to the systems-scale workflows now standard in genomics and bulk proteomics. Large-scale interactome screens extended quantitative AE-MS to thousands of baits but required years of sample preparation, milligram-scale input per bait, and a stable cell line generated for each construct^12,13,16^. Extending such efforts to dynamic, condition-resolved interactomes, or to sample-limited systems such as endogenously expressed receptors, has therefore remained largely out of reach. This same limitation constrains the experimental validation of the rapidly expanding sets of computationally predicted and designed protein interactions^31–33^.

HIP-MS changes what is routinely feasible in interactomics along three axes. The first is **time**. Condition- and time-resolved screens that previously demanded dedicated multi-month efforts can now be performed in days, with all conditions and replicates measured side-by-side on a single plate using the same reagents, handling, and MS acquisition batch. This eliminates the inter-experiment variability that has long limited the interpretability of dynamic AE-MS data and yields timepoint measurements that are directly and quantitatively comparable. Our TNFR1 and rapamycin experiments illustrate this directly: the temporal assembly of Complex I and the kinetically separable ESCRT-III arm, and stage-resolved autophagy interactions across many baits in parallel, reflect an experimental design effectively inaccessible at conventional throughput **(Fig. 5).** The second axis is **sample input**. Because on-plate capture eliminates bead handling and the sample losses of repeated transfer steps, specific enrichment no longer requires the milligram-scale inputs of conventional AE-MS. In our EMC2 titration, the near-complete EMC was recovered from 10 µg, four subunits remained enriched at 1.25 µg, and the bait itself was robustly captured from just 312 ng of lysate **(Fig. 4a)**. This brings within reach interactomes from intrinsically scarce material such as sorted primary populations, organoids, and low-abundance or membrane-resident proteins. The endogenous TNFR1 experiment, in which an unmodified receptor was pulled down through a tagged extracellular ligand, shows that low-input capture and endogenous targeting can be combined in a single 96-well experiment **(Fig. 5)**. The third axis is **combinatorial scale**. Because the per-experiment overhead of plate-based HIP-MS is largely fixed, the marginal cost of adding one more condition, timepoint, or bait is small: scaling from a few dozen pulldowns to a full plate of 384 increases the experimental cost only modestly, while expanding the biological scope of the experiment. This changes the economics of interactomics and makes exhaustive screens across drug treatments, genetic modifications, cell lines, or environmental conditions tractable in a single batch, rather than requiring incremental design and execution over months. Together with built-in replicates and shared-background normalization across thousands of pulldowns, this brings AE-MS into a regime where systems-level interactomics becomes feasible.

These gains come with trade-offs that define the current operating envelope of the platform. First, like other ORF-based screens, HIP-MS expresses baits exogenously by transient transfection, with the attendant caveats of non-native abundance and the potential for stoichiometry- or localization-dependent artifacts; the endogenous TNFR1 capture indicates one route around this, but the default bait mode remains overexpression. Second, the throughput that defines the platform comes at a modest cost in depth and in the recovery of the weakest interactions: the 500 SPD gradient quantifies fewer proteins than the 100 SPD gradient, and while it returns a higher proportion of bona-fide interactors, the longer gradient extends detection into the substoichiometric range. Finally, precision and recall are bounded by incomplete interaction annotation, and individual non-reciprocal observations (e.g., ATG4B–WIPI2, **Fig. 5d**) remain candidate findings pending orthogonal validation. Each of these is addressable, e.g., through longer gradients or on-plate cross-linking for weak interactions, and multi-subunit baiting to raise per-complex recovery.

A defining feature of HIP-MS is that it is modular. Each of the five components introduced above (the affinity tag, the bait expression strategy, the culture and lysis format, the on-plate capture chemistry, and the LC-MS acquisition) can be swapped or upgraded independently, so the platform is not tied to any single reagent or instrument. This flexibility also makes it extensible. We chose the ALFA system for its favorable combination of affinity, tag size, and low background, but the on-plate format is in principle compatible with any immobilized affinity reagent, including conventional FLAG, HA, or GFP systems, allowing existing bait libraries to be ported into HIP-MS without re-cloning. The ALFA system itself can be diversified: NbALFA variants with tunable affinities could enable on-plate elution by peptide competition, opening downstream applications such as native MS on the same plate^24^. The capture step could be combined with crosslinking to stabilize transient interactions before washing, while cell culture, transfection, and lysis (currently in 96 well format) could be compressed into 384-well to match the downstream capture geometry. Lysates generated for HIP-MS pulldowns are also directly compatible with orthogonal readouts for interactions such as PELSA, which maps binding interfaces at peptide-level resolution and can independently validate the pulled-down complexes^34^.

Interactome biology increasingly depends on the integration of experimental and computational protein science, and HIP-MS is well-suited to operate at this interface^35,36^. Co-folding methods such as AlphaFold-Multimer, AlphaFold3, RoseTTAFold, OpenFold, and Boltz^31,37–40^ and de novo protein design pipelines^33,41,42^ are now generating predicted interactions and engineered binders at a rate that far outpaces conventional experimental validation. HIP–MS can absorb the throughput this will require: predicted or designed interactions can be expressed as ALFA-tagged constructs and tested in parallel for binding specificity, off-target interactions, and context-dependent behavior. Just as importantly, the platform can supply what current predictors most lack on the input side. Their performance is weakest for transient, condition-dependent, and cross-species interactions, with the sharpest drop at pathogen–host interfaces^9,15^, where evolutionary distance from the training distribution leaves little signal to learn from. Closing this gap will require broadening the training distribution itself with large-scale experimental interaction data, rather than only refining model architecture. This is exactly the kind of data HIP-MS is built to generate, across conditions, perturbations, and cell types.

## Methods

### Cell culture

HEK293F cells were maintained in suspension in FreeStyle 293 expression medium [Gibco, 12338-018] at 37 °C and 8% CO₂ on an orbital shaker at 120 r.p.m. between 0.5 and 4 × 10⁶ cells/mL. HEK293T and A549 cells were maintained in DMEM (high glucose, Capricorn Scientific, [DMEM-HPSTA]) supplemented with 10% FBS and 1% penicillin–streptomycin [Gibco, 15140-122] at 37 °C and 5% CO₂, grown in 10- or 15-cm dishes [Sarstedt, 83.3902 and 83.3903] and passaged at 80–90% confluency with 0.05% trypsin–EDTA.

### Plasmid cloning

ALFA-tagged human ORFs were either ordered from IDT or cloned with an N-or C-terminal ALFA tag^24^ from cDNA **(Supplementary table 1)**. All transfections were performed in 96-well format with four transfection replicates per construct unless stated otherwise.

### Benchmarking plasmids in suspension HEK293F

A 96-well DNA plate of 50 ORFs spanning different cellular compartments and controls (100 ng/µL) was used to transfect four 96-deep-well plates. Of the 50 ORFs in our benchmark panel, we curated 25 baits for the per-bait display panels, applying three inclusion criteria: (i) the bait protein itself self-enriched (FDR < 0.05, log₂ FC > 1.5); (ii) at least one annotated interactor existed in STRING (combined score > 700) or Complex Portal; and (iii) ≤ 150 significantly enriched proteins overall, to exclude promiscuous baits whose enriched sets are dominated by off-target signal (e.g., SRSF1 with 1,154 enriched proteins, HMGB1 with 1,098, COPB2 with 316). Baits that did not self-detect (e.g., LC3, RPL38, RPS10) or had no STRING/Complex Portal annotation were also excluded. Each well received 250 ng plasmid DNA and 1 µL linear polyethylenimine (PEI STAR, TOCRIS, [8174], 1 mg/mL) diluted in 50 µL Opti-MEM [Gibco, 31985-062], respectively. Complexes were mixed, incubated for 20–30 min, and 50 µL was dispensed per well into 96-deep-well plates [Corning, 3960] containing 500 µL of HEK293F cells at 2 × 10⁶ cells/mL. Cells were cultured for 48 h prior to lysis.

### Benchmarking plasmids in adherent HEK293T

HEK293T cells were seeded into 96-well plates [Sarstedt, 83.3924.005] at 40,000 cells/well in 90 µL and grown for 24 h to ∼50% confluency. The same 50 baits were used as described above (100 ng/µL). Per well, 1 µL of plasmid in 4 µL of Opti-MEM was combined after 5 min incubation with 0.2 µL of Lipofectamine 2000 [Invitrogen, 11668019] diluted with 5 µL Opti-MEM solution per well; complexes were formed for 30 min and 10 µL was added per well with the INTEGRA Mini 96 [Integra Biosciences, 4801]. Six 96-well plates (four replicates per three 96-well plates) were transfected and lysed in two different buffers (three plates each; see “Cell lysis”).

### EMC2-ALFA and GFP for the lysate titration experiment

HEK293T cells were seeded at 12 × 10⁶ cells per 15-cm dish and grown for 24 h. One dish was transfected with ALFA-tagged EMC2 and one with ALFA-GFP using the same Lipofectamine 2000 chemistry. 200 µL of plasmid (20 µg) in 800 µL of Opti-MEM was combined with 40 µL of Lipofectamine 2000 in 1 mL of Opti-MEM, complexes formed for 30 min, and 2 mL added dropwise to each dish (18 mL of medium). Cells were cultured for 24 h prior to lysis.

### Autophagy plasmids in adherent HEK293T

A panel of 18 ALFA-tagged autophagy-related ORFs plus five autophagy constructs from the benchmarking panel (LC3, Beclin-1, FKBP1A, MLST8, DYNLL1; Supplementary Table 1) was arrayed at 100 ng/µL. HEK293T cells were seeded into 96-well plates at 20,000 cells/well in 90 µL of complete medium and grown for 30 h. Transfection followed the benchmarking protocol but with FuGENE 4K [Promega, E5911] in place of Lipofectamine 2000.

### Cell treatment

#### Rapamycin time-course

48 h after transfection (autophagy pulldowns) or seeding (rapamycin µPhos), HEK293T cells were treated with 200 nM rapamycin in 0.1% DMSO [Sigma Aldrich, D2650-100ML] for 30 min, 1 h, 2 h or 4 h, with matched 0.1% DMSO vehicle controls. Rapamycin was diluted from a 20 mM DMSO stock to 2,000 nM in 1% DMSO/DMEM; 10 µL was added per well (100 µL culture medium) with an INTEGRA 12-channel pipette, and vehicle wells received 10 µL of 1% DMSO/DMEM. Treatments were started from the longest time point so all plates reached lysis simultaneously.

#### TNF time-course

Recombinant ALFA-tagged TNF was designed and purified by our in-house protein production facility (ALFA-IZ-TNFα; design based on isoleucine-zipper-stabilized TNF-superfamily ligands^43,44^, applied here to TNF; 1 mg/mL in PBS + 0.1% BSA, provided by the protein production core facility). Recombinant TNF was diluted to 0.5 ng/µL in pre-warmed complete medium. For 96-well A549 plates, medium was replaced with 100 µL of TNF-containing medium (50 ng/well) in a staggered manner so the 10-, 5-, 2- and 1-min time points ended simultaneously, with an untreated cell control and a medium control. For 10-cm dishes, 3 mL of medium was replaced (1.5 µg/dish) for 15 min at 37 °C; for 15-cm dishes (TNF phosphoproteomics), 15 mL was replaced (7.5 µg/dish) for 15 min at 37 °C. Untreated controls were processed in parallel; treatments were stopped on dry ice and TNF-containing medium removed.

### Cell lysis

#### Lysis for affinity enrichment

Wash buffer was 50 mM Tris-HCl [Sigma Aldrich, 0000303827] pH 7.5 at 4 °C, 150 mM NaCl [Sigma Aldrich, 0000395932], 5% glycerol [Thermo Scientific, 158920025]. IP lysis buffer was wash buffer + 0.5% IGEPAL CA-630 [Sigma-Aldrich, 18896-50ML]. GentleLys buffer was bought from Cube Biotech [Cube Biotech, 18907]. Lysis buffers were freshly supplemented with cOmplete EDTA-free protease inhibitor (one tablet per 50 mL, Roche, [05056489001]), 1 mM MgCl₂ and 0.1% (v/v) in-house benzonase.

##### HEK293F (benchmarking)

at 48 h post-transfection, deep-well plates were centrifuged at 1,000g for 3 min, medium decanted by plate inversion, and cells lysed in 250 µL/well of IP lysis buffer on ice with the INTEGRA Mini 96, shaken at 950 r.p.m. for 30 min at 4 °C and cleared at 4,000g for 15 min at 4 °C.

##### HEK293T (benchmarking and autophagy)

cells were lysed on-plate in 100 µL/well of ice-cold lysis buffer with the INTEGRA Mini 96 (benchmarking: three of six 96-well plates with GentleLys, three with IP lysis buffer; autophagy: GentleLys only), mixed by pipetting 30 cycles with 60 µL, shaking at 950 r.p.m. for 120 min at 4 °C, transferred to Eppendorf Twin.tec 96-well plates [Eppendorf, 0030 129.512] with an INTEGRA 1250 pipette, and cleared at 4,000g for 15 min at 4 °C. **HEK293T (titration):** cells were washed twice with 6 mL of PBS [Capricorn, PBS-1A], lysed in 200 µL of IP lysis buffer per 15-cm dish, scraped into 1.5-mL LoBind tubes, rotated for 45 min at 4 °C, and cleared at 18,000g for 15 min at 4 °C.

##### A549 (96-well TNF pulldown)

after dry-ice stop, cells were washed once with 100 µL/well of ice-cold TBS pH 7.4 (50 mM Tris-HCl, 150 mM NaCl) and lysed in 100 µL/well of GentleLys lysis buffer; cells were detached by pipetting 10 times, shaken at 700 r.p.m. for 1 h at 4 °C, transferred to Twin.tec LoBind 96-well plates, and cleared at 10,000g for 15 min at 4 °C. **A549 (10-cm dish bead pulldown):** cells were washed three times with 5 mL of PBS and lysed in 2 mL of GentleLys lysis buffer per dish; lysates were scraped into 2-mL LoBind tubes, shaken at 1,200 r.p.m. for 1.5 h at 4 °C, and cleared at 18,000g for 15 min at 4 °C.

#### Protein quantification (titration experiment)

Cleared lysates were quantified by Pierce BCA Protein Assay [Thermo Fisher Scientific, 23225] in triplicate against a BSA standard curve and measured on a Tecan plate reader [Tecan Infinite M Plex, 30050303]. HEK293T lysate was diluted to 2 mg/mL and HEK293F lysate to 4 mg/mL in IP lysis buffer. Remaining material was snap-frozen and stored at –20 °C.

#### SDC lysis (µPhos)

SDC lysis buffer was 2% sodium deoxycholate [Roth, CN30.3], 50 mM TEAB [Merck, 18597-100ML], 10 mM TCEP, 40 mM 2-chloroacetamide, pH 8.5. **Rapamycin µPhos (96-well HEK293T):** on ice, medium aspirated without washing, 19 µL of SDC lysis buffer added per well, shaken 800 r.p.m. for 1 min, pulse-spun, heated 75 °C and 1,200 r.p.m. for 10 min in an Eppendorf ThermoMixer (reduced temperature for the polystyrene plate), sonicated 10 min in a water-bath sonicator (10 s on / 10 s off, 4 °C), pulse-spun. **TNF µPhos (15-cm dishes):** after dry-ice stop, cells were washed three times with ice-cold TBS pH 7.4 (to remove serum albumin), lysed in 500 µL of SDC lysis buffer per dish, scraped into 1.5-mL LoBind tubes, heated 95 °C and 1,200 r.p.m. for 10 min, sonicated 30 s with a probe sonicator (20%, output 3), cooled, heated again 5 min at 95 °C, cleared at 18,000g for 10 min. Protein was quantified by tryptophan fluorescence (100 µL of 8 M urea/50 mM TEAB per well; standards 0–25 ng/µL in duplicate; sample at 1×, 1:2, 1:5, 1:10). 20-µg aliquots were snap-frozen and stored at –20 °C.

### Affinity enrichment

#### Anti-ALFA nanobody plate coating

##### 384-well format

streptavidin-coated 384-well plates [Thermo Scientific, 15405] were washed three times with 100 µL/well of wash buffer on an automated plate washer [Cytena C. Wash Plus, CYT-CWP-2024-01-016]. Biotinylated anti-ALFA nanobody (commercial, NanoTag, [N1505-Biotin], or in-house produced) was diluted to 2.5 ng/µL and dispensed at 50 µL/well. Plates were incubated for 2 h at 25 °C and 1,200 r.p.m., washed three times with wash buffer, sealed and stored at 4 °C. **96-well format (TNF receptor pulldown):** pre-blocked streptavidin-coated 96-well plates [Thermo Scientific, 15124] were coated with 50 µL/well of biotinylated anti-ALFA nanobody at 0.5 µg/well in PBS for 2 h at room temperature and 900 r.p.m., and washed three times with wash buffer.

#### On-plate affinity enrichment and washes

Cleared lysate was transferred to the nanobody-coated plate at 50 µL/well (suspension benchmarking, titration), 90 µL/well (adherent benchmarking, autophagy), or 100 µL/well (96-well TNF pulldown). For HEK293F benchmarking, the remaining 200 µL/well was archived as snap-frozen aliquots in a heat-sealed 96-deep-well LoBind plate at –20 °C. Plates were sealed and incubated 12–17 h at 4 °C and 1,200 r.p.m. (3 h at 700 r.p.m. for the TNF 96-well pulldown), briefly centrifuged, and washed three times with 100 µL/well of ice-cold wash buffer (+0.01% DDM [Roth, CN26.3] where indicated) on the automated plate washer.

#### Bead-based affinity enrichment (10-cm dish comparator)

25 µL of ALFA Selector ST magnetic agarose bead slurry (50%; NanoTag, N1516) was equilibrated by three washes with 1 mL of wash buffer on a magnetic rack on ice. Cleared lysate from one 10-cm dish was split into three aliquots (technical replicates) and incubated overnight with beads at 4 °C and 1,200 r.p.m. Beads were washed three times with 1 mL of wash buffer.

#### On-plate digestion and quenching

Digestion buffers were prepared fresh. Buffer 1 (6 M urea [Sigma Aldrich, U1250-1KG], 10 mM Tris-HCl pH 8.5, 3 mM DTT, 1 ng/µL LysC) was dispensed at 20 µL/well and incubated 3 h at 30 °C and 1,200 r.p.m. Buffer 2 (10 mM Tris-HCl pH 8.5, 0.4 ng/µL Promega Trypsin Gold MS-grade [Promega, V5111/VA900A], 7.5 mM 2-chloroacetamide) was added at 40 µL/well; digestion continued at 37 °C and 1,200 r.p.m. for 5 h (HEK293T benchmarking, titration) or overnight (HEK293F benchmarking, autophagy). For membrane-protein experiments (benchmarking, titration, autophagy), both digestion buffers and the wash buffer included 0.01% DDM. Reactions were quenched with 6.6 µL/well of 10% TFA [Sigma Aldrich, T0699-100ML]. The TNF 96-well pulldown used higher enzyme loads in larger volumes — 50 µL of buffer 1 with 100 ng LysC/well (3 h, 30 °C, 1,000 r.p.m.); 100 µL of buffer 2 with 200 ng trypsin/well (overnight, 37 °C, 800 r.p.m.); quench 15 µL of 10% TFA.

#### On-bead digestion (10-cm dish comparator)

Beads were digested in 50 µL of buffer 1 (6 M urea, 10 mM Tris-HCl pH 8.5, 3 mM DTT, 100 ng LysC, 0.01% DDM) for 3 h at 30 °C and 1,200 r.p.m., followed by 100 µL of buffer 2 (10 mM Tris-HCl pH 8.5, 200 ng Promega trypsin, 7.5 mM CAA, 0.01% DDM) for 4.5 h at 37 °C and 1,200 r.p.m. Beads were spun, supernatants transferred to fresh tubes, and digests quenched with 15 µL of 10% TFA.

### Sample preparation for LC-MS analysis

#### Evotip loading (affinity-enriched samples)

Evotips [Evosep Biosystems, EV2011] were activated with 50 µL of buffer B (acetonitrile, 0.1% FA) at 700g for 1 min, soaked in 2-propanol for 1 min, and washed with 50 µL of buffer A (water, 0.1% FA) at 700g for 1 min. Acidified peptides were loaded at 500g for 2–3 min and washed with 250 µL of buffer A at 400g for 1 min. Load volumes were 30 µL (HEK293F and HEK293T benchmarking; two 30-µL aliquots on separate Evotips for the 500 and 100 SPD analyses of HEK293T), the entire ∼66 µL sample (titration, autophagy), 50 µL (TNF 96-well) or 75 µL (TNF beads). Tips were stored at 4 °C in buffer A until LC-MS.

#### Phosphopeptide enrichment (µPhos)

µPhos buffers were prepared as described^45^. **Rapamycin µPhos:** 2 µL of digestion buffer was added per well, plates sealed with a silicone mat and digested 3 h at 37 °C and 1,200 r.p.m. **TNF µPhos:** 5 µg of protein per replicate in 19 µL of 2% SDC lysis buffer was digested with 1 µL of digestion buffer for 4 h at 37 °C and 1,500 r.p.m. Enrichment was performed as previously described^45^ and samples were loaded onto pre-equilibrated Evotips (washed once with 50 µL of buffer B, activated for 1 min in 2-propanol, equilibrated with 50 µL of buffer A). All Evotip centrifugation steps were at 700g for 1 min, except sample loading (4 min). Tips were stored at 4 °C in buffer A until LC-MS.

#### MS acquisition

Data were acquired on an Evosep Eno coupled to an Orbitrap Astral Zoom or Orbitrap Astral. 500 SPD measurements (HEK293F and HEK293T benchmark panels, EMC2 titration, rapamycin time-course on the Astral Zoom) used the Evosep Performance column (4cm × 150 µm, 1.9 µm particles); 100 SPD measurements (HEK293F benchmark on the Astral Zoom; HEK293T benchmark and TNF-receptor pulldowns on the Astral) used the column for 100/60 SPD (8 cm × 150 µm, 1.5 µm particles); both with a 30-µm stainless-steel emitter in the Evosep bullet adapter. The EASY spray source was operated at 2,000 V with FAIMS at –40 V (inner 100 °C, outer 80 °C). The DIA method comprised an MS1 scan from 380–980 m/z at 240k resolution (3 ms IT, 500% AGC) and MS2 scans in the Astral analyzer (500% AGC; 5 ms IT for 500 SPD, 6 ms for 100 SPD) using 100 variable windows (500 SPD) or 3 Th fixed windows (100 SPD) across 380–980 m/z with an NCE of 25% for HCD fragmentation; windows were designed with pydiaid^46^ from a reference HeLa proteome. Phosphoproteomics data were acquired on the Orbitrap Astral Zoom using an IonOpticks Aurora Rapid column (75 µm × 5 cm, 1.7 µm) and the Whisper Zoom 120 gradient; spray voltage 1,900 V, MS1 380–1,380 m/z, 100 variable MS2 windows over the same range, 10 ms IT.

### Data analysis

#### Library prediction and DIANN search

Spectral libraries were predicted with DIA-NN from human protein sequences using tryptic digestion with fixed cysteine carbamidomethylation, variable methionine oxidation, and up to two variable modifications. The benchmarking and EMC2-titration library (DIA-NN 2.3.1) was built from UniProt human SwissProt augmented with NbALFA, ALFA-tag, streptavidin, GFP, BSA, LysC and trypsin sequences; the rapamycin library (DIA-NN 2.3.1) from plain human SwissProt; and the TNF-receptor library (DIA-NN 2.2.0) from the UniProt human reference proteome, with protein N-terminal acetylation additionally enabled for the latter two. Raw files were searched in DIA-NN 2.1.0 with peptidoform scoring and match-between-runs at 7 ppm (MS1) / 10 ppm (MS2) mass tolerances (15 ppm for the TNF-receptor pulldowns)^47^.

#### Protein quantification and differential enrichment

Protein abundances were quantified per condition by two methods: MaxLFQ from the DIA-NN protein-group table and directLFQ^48^ (version 0.3.3) from proteotypic precursors (Q value < 0.01); both were log₂-transformed. Missing values were imputed with Gaussian draws from N (μ − 1.8·σ, 0.3·σ), where μ and σ are the mean and standard deviation of the observed log₂ values (random seed = 42). Differential enrichment was performed for each cell-line × gradient combination on the imputed directLFQ values: each bait gene (3–10 replicate wells; “none” and “GFP” controls excluded as targets) was contrasted against all other samples, including the control wells, using an empirical-Bayes moderated t-test (alphapepttools 0.2.0^49^). P-values were corrected to FDR by Benjamini–Hochberg, and proteins with FDR < 0.05 and log₂ fold-change > 1.5 were called significantly enriched. For the rapamycin/autophagy time-course (Fig. 5d, Supplementary Fig. 5g), samples were first restricted to those in which the bait protein ranked among the 50 most intense protein groups (MaxLFQ), removing wells with weak bait signal. Partner-intensity trajectories were computed for six bait-partner pairs at three timepoints (30 min, 1 h, 4 h; the 2 h plate failed quality control and was excluded) as the difference between the median MaxLFQ log₂ intensity of the partner in rapamycin versus matched DMSO wells, with undetected values set to the 1st-percentile floor before differencing. For the accompanying significance view (**Supplementary Fig. 5g**), partner enrichment was tested separately in the rapamycin and DMSO pulldowns of each bait against a common background of the other baits on the plate (excluding baits of the same gene or autophagy stage), using the alphapepttools eBayes test described above.

#### Curation of TNF and mTOR/autophagy pathway phosphosites

Phosphosite panels were compiled from canonical literature and PhosphoSitePlus to span major arms of each cascade, prioritizing kinase activation-loop residues and well-established substrate sites^50–54^. Where the same residue was detected at multiple multiplicities, the peptidoform with the lowest adjusted p-value was selected. Each retained site reached |log₂FC| ≥ 0.5 in the expected direction at ≥ 1 time point; per-cell asterisks in **Supp. Fig. 5e** mark cells passing FDR < 0.05.

## Author contributions

L.G. and E.Z. designed and developed the HIP-MS platform, performed experiments, analyzed data, and wrote the manuscript. T.H. developed the mass spectrometry methods used in the paper and helped plan experiments. C.P. contributed to experiments and method development. V.B. contributed to data analysis and accessibility. A.C.M. helped to design the pipeline and supervised the work. M.M. conceived and supervised the project. All authors edited and approved the final manuscript.

---

**Supplementary Figure 1.**
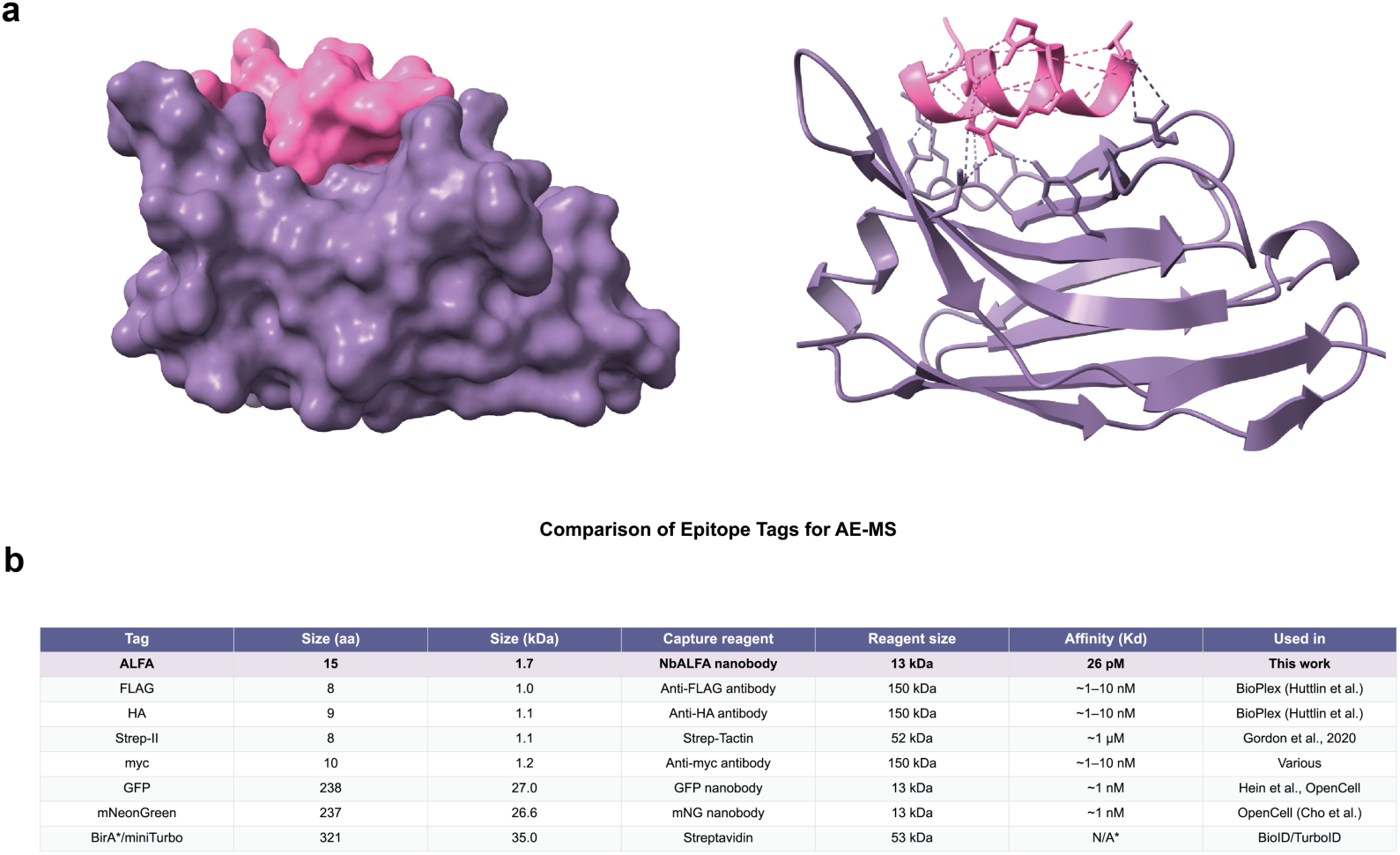
The ALFA-tag / NbALFA capture system: structural basis and size comparison with conventional affinity reagents. (a) AlphaFold3 predicted structure of the NbALFA single-domain antibody (purple) bound to the 15-amino-acid ALFA-tag peptide (pink). Left: surface map, right: secondary structure representation of the same complex^24^. (b) Size and property comparison of common epitope tag and affinity-capture reagents used in published interactome workflows.

**Supplementary Figure 2.**
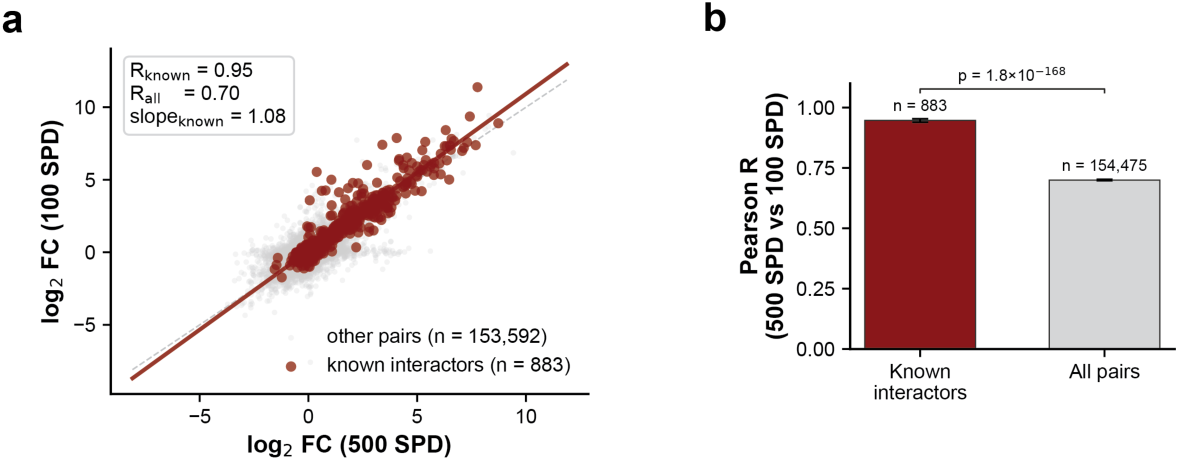
Cross-gradient reproducibility on the full bait–prey set. (a) Side-by-side log fold-change scatter, 500 SPD vs 100 SPD, for all bait-prey pairs with finite fold change in both gradients (n = 153,592 pairs; grey, light dots) overlaid with known interactors (n = 883 pairs; dark red dots). Solid line = least-squares fit on the known subset; dashed line = unity. Pearson R (two-sided). (b) Pearson correlation coefficients with 95% confidence intervals (Fisher z-transformation) for the known-interactor and all-pair subsets, with a two-sided two-sample Fisher z-test for the difference between correlations, error bars: 95% CI; sample sizes shown above each bar.

**Supplementary Figure 3.**
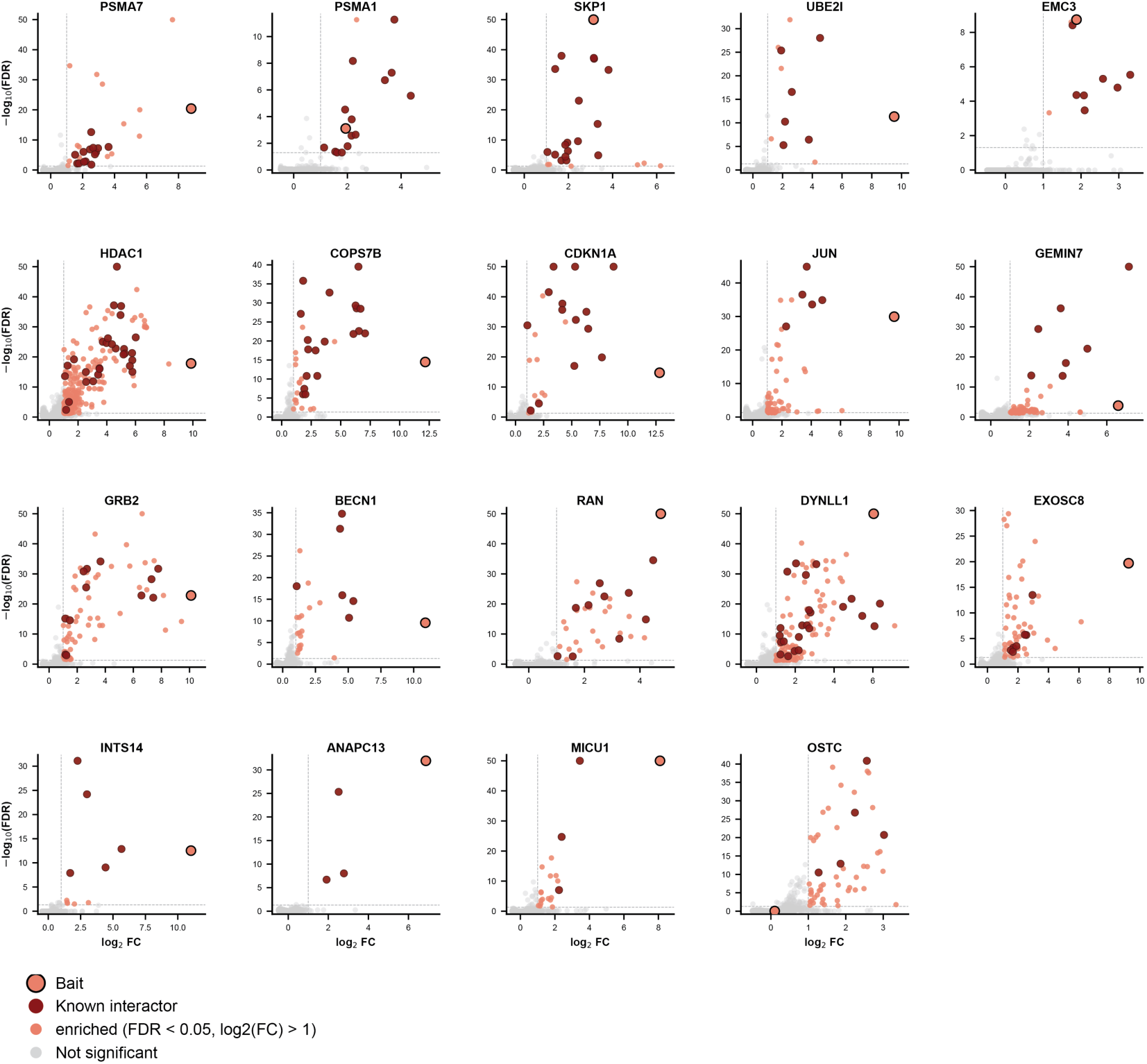
Per-bait volcano plots for 19 curated baits. Remaining 19 curated baits for the pulldowns in HEK293F cells at 500 SPD not shown in Fig.3a. Differential enrichment was done with empirical-Bayes moderated t-test on directLFQ-Gaussian-imputed data (two-sided p-values, Benjamini–Hochberg FDR).

**Supplementary Figure 4.**
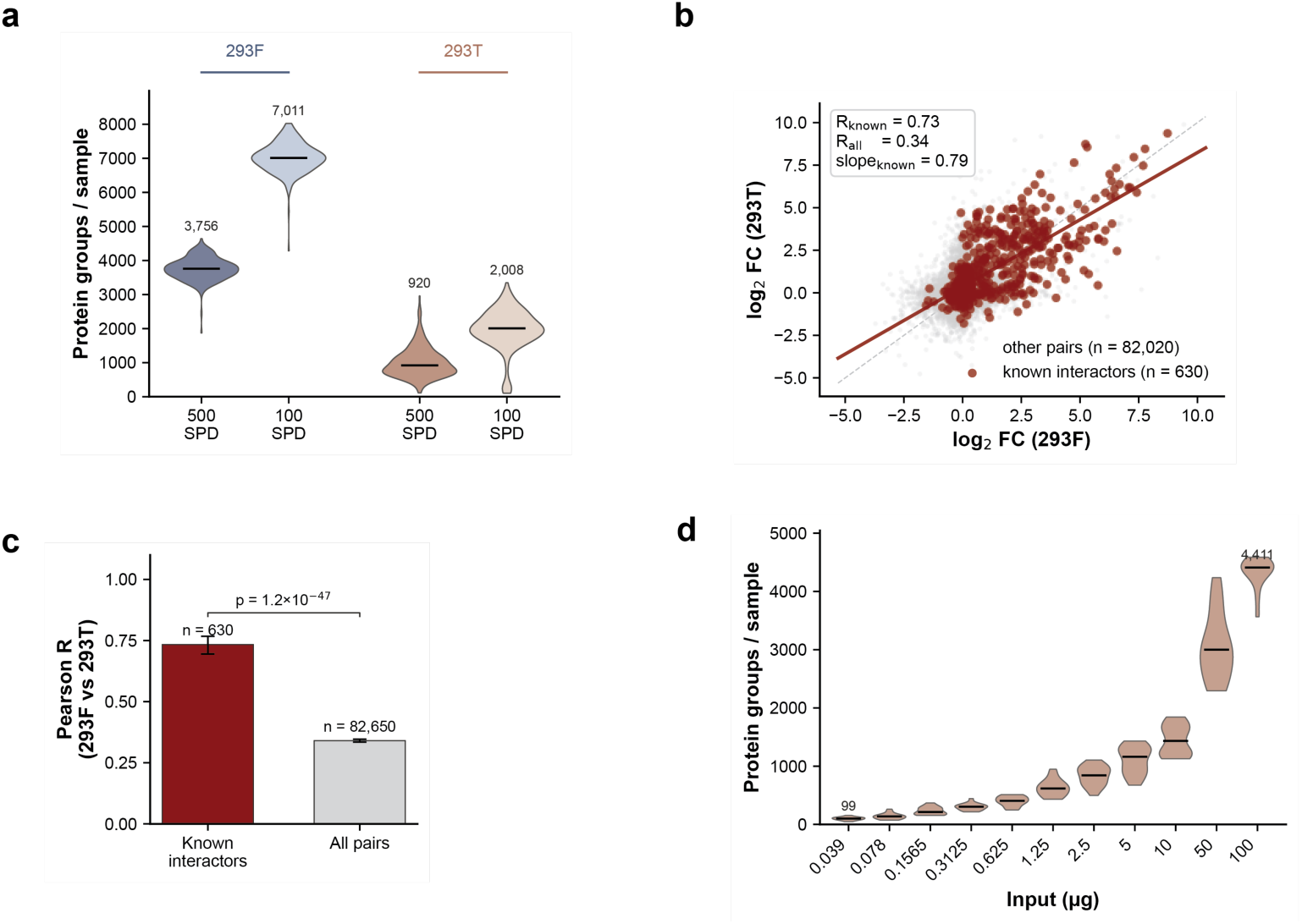
Cross-cell-line depth and reproducibility context for Figure 4. (a) Per-sample protein identifications across the four arms (293F 500 SPD, 293F 100 SPD, 293T 500 SPD, 293T 100 SPD). (b) All bait–prey pairs scatter across cell lines at 500 SPD. Same axes as main Fig. 4c but extends the comparison from known interactors only to every bait-prey pair with a finite log fold change in both cell lines. Aggregated across the 25 curated baits (n = 4 wells per bait per cell line). (c) Pearson correlation coefficients for the all-pair and known-interactor subsets, with 95% confidence intervals (Fisher z-transformation). Two-sided two-sample Fisher z-test comparing the two correlations. (d) Per-sample proteins quantified across the 11 EMC2-titration amounts (293T, 500 SPD). One violin per concentration; each dot is one sample (EMC2 and GFP wells pooled at each level)

**Supplementary Figure 5.**
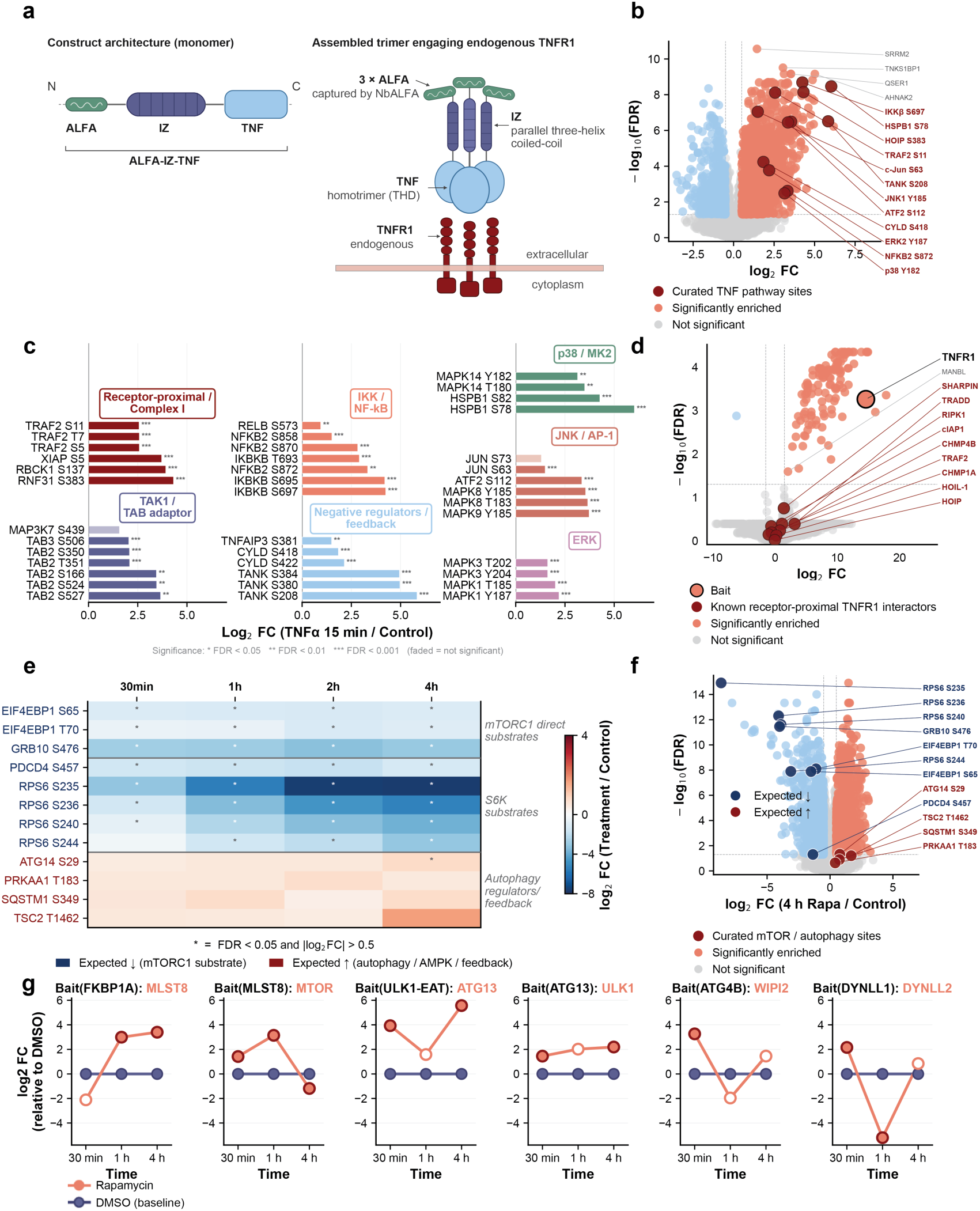
Orthogonal phosphoproteomic and bead-based AE-MS validation of TNF and rapamycin signaling. (a) Architecture of the ALFA-IZ-TNFα construct used for endogenous TNFR1 capture in Fig. 5a–c. (b) Volcano plot of the global phosphoproteome response to TNFα (15 min, 100 ng/ml) vs. unstimulated control in A549 cells measured at 120 SPD; n = 7 biological replicates per condition. Dark red: twelve curated TNF pathway sites spanning Complex I, LUBAC, IKK, NF-κB, p38/MK2, JNK/AP-1, ERK, and feedback regulators^50,51^, labeled regardless of significance. Grey: top four non curated up-regulated sites. Salmon: significantly enriched; pastel blue: significantly depleted (FDR < 0.05, |log FC| > 0.5). (c) Forest plot of the twelve curated TNF pathway sites from (b) and additional pathway members, grouped by signaling subgroup (Receptor-proximal/Complex I, TAK1/TAB adaptor, IKK/NF-κB, Negative regulators/feedback, p38/MK2, JNK/AP-1, ERK), shown across three columns corresponding to upstream → NF-κB axis → MAPK cascades. Bar length = log FC vs. control; colors match subgroup headers. Faded bars: not significant. Asterisks: *FDR < 0.05, **<0.01, ***<0.001. (d) Volcano plot of ALFA-TNF pulldown comparing 15 min TNFα stimulation vs. unstimulated control, performed with ALFA Selector ST magnetic agarose beads from 10-cm dishes (A549, 500 SPD; n = 3 biological replicates). Large bordered circle: bait (TNF, FDR = 0.05, |log FC| = 1). (e) Heatmap of twelve curated mTORC1/autophagy phosphosites across a rapamycin time-course (30 min, 1 h, 2 h, 4 h; HEK293T, µPhos, 120 SPD; n = 12 per timepoint), grouped into mTORC1 direct substrates, S6K substrates, and autophagy/feedback regulators^50,52–54^. Asterisks: FDR < 0.05, |log FC| > 0.5. (f) Volcano plot of the global 4 h rapamycin vs. control phosphoproteome (48,814 sites). Curated sites from (e) overlaid in navy (expected ↓) and dark red (expected ↑). Coloring of all other points as in (b). (g) Statistical view of the bait-partner trajectories in Fig. 5d. For each bait and timepoint, partner enrichment was tested separately in the rapamycin and the DMSO pulldown against a common background of the other baits on the plate (excluding baits of the same gene or autophagy stage), using empirical-Bayes moderated t-test on Gaussian-imputed log₂ intensities. Plotted is the difference between the two enrichment log₂ fold-changes (rapamycin − DMSO), with the DMSO baseline at zero. Filled circles mark a robustly enriched partner with a rapamycin − DMSO enrichment difference > 1 log₂ unit (Fisher-combined enrichment p < 0.05).

